# Exhaustively Identifying Cross-Linked Peptides with a Linear Computational Complexity

**DOI:** 10.1101/097089

**Authors:** Fengchao Yu, Ning Li, Weichuan Yu

## Abstract

Chemical cross-linking coupled with mass spectrometry is a powerful tool to study protein-protein interactions and protein conformations. Two linked peptides are ionized and fragmented to produce a tandem mass spectrum. In such an experiment, a tandem mass spectrum contains ions from two peptides. The peptide identification problem becomes a peptide-peptide pair identification problem. Currently, most existing tools don’t search all possible pairs due to the quadratic time complexity. Consequently, a significant percentage of linked peptides are missed. In our earlier work, we developed a tool named ECL to search all pairs of peptides exhaustively. While ECL does not miss any linked peptides, it is very slow due to the quadratic computational complexity, especially when the database is large. Furthermore, ECL uses a score function without statistical calibration, while researchers^1,2^ have demonstrated that using a statistical calibrated score function can achieve a higher sensitivity than using an uncalibrated one.

Here, we propose an advanced version of ECL, named ECL 2.0. It achieves a linear time and space complexity by taking advantage of the additive property of a score function. It can analyze a typical data set containing tens of thousands of spectra using a large-scale database containing thousands of proteins in a few hours. Comparison with other five state-of-the-art tools shows that ECL 2.0 is much faster than pLink, StavroX, ProteinProspector, and ECL. Kojak is the only one tool that is faster than ECL 2.0. But Kojak does not exhaustively search all possible peptide pairs. We also adopt an *e*-value estimation method to calibrate the original score. Comparison shows that ECL 2.0 has the highest sensitivity among the state-of-the-art tools. The experiment using a large-scale *in vivo* cross-linking data set demonstrates that ECL 2.0 is the only tool that can find PSMs passing the false discovery rate threshold. The result illustrates that exhaustive search and well calibrated score function are useful to find PSMs from a huge search space.

## 1 Introduction

The power of chemical cross-linking coupled with mass spectrometry (XL-MS) has been well demonstrated in understanding protein structures and protein-protein interactions ^3-6^.

In XL-MS, we first link proteins with a cross-linker. Then, we quench the reaction and digest the proteins. Finally, *we* obtain pairs of linked peptides. However, identifying cross-linked peptides from XL-MS data is computationally challenging. The time complexity is quadratic with respect to the number of peptides in the database. Consequently, exhaustively searching all peptide-peptide pairs is time consuming and resource demanding. For example, there are around 3 × 10^6^ peptides in the Homo sapiens (human) database (UniProtKB / Swiss-Prot, 2015-11 release, 20,205 proteins). Suppose the precursor mass tolerance is 10 ppm (parts per million). There will be on average around 10^7^ peptide-peptide candidate pairs for each experimental spectrum.

Many methods^7-27^ have been developed to identify cross-linked peptides. These methods can be classified into two groups. The first group converts searching peptide-peptide pairs into searching two peptides sequentially with the help of specific cross-linkers. The second group limits the number of peptide-peptide pairs with heuristic pre-filtering procedures.

Methods in the first group convert the quadratic time complexity into a linear time complexity by using cross-linkers^28-31^that can be broken during dissociation (e.g. collision-induced dissociation (CID)). Kaake et al.^31^ and Kao et al.^29^ proposed to couple such cross-linkers with three levels of mass spectrometry (i.e. MS1, MS2, and MS3). The issue is that generating three levels of mass spectra requires a longer cycle time. Liu et al.^32^ and Götze et al.^8^ proposed to use cross-linker-cleaved signature peaks to infer the masses of two peptides. This method avoids generating three levels of mass spectrometry. However, the signature peaks may not be observed all the time, resulting in loss of useful data. Furthermore, the cleavable cross-linkers are not as widely used as mass-spectrometry-noncleavable cross-linkers (such as disuccinimidyl suberate (DSS) and bis(sulfosuccinimidyl) suberate (BS3)) in biological experiments^33,34^.

Methods in the second group include xQuest/xProphet^35,36^, pLink^23^, ProteinProspector^25^, StavroX^7^, and Kojak^27^. They only keep a fixed number of peptides to generate peptide-peptide pairs for each experimental spectrum. For example, xQuest/xProphet first uses the top 5000 peptides for pairing. Then, it Liters all peptide-peptide pairs with a pre-score. Finally, it uses the top 50 peptide-peptide pairs for fine scoring. Similarly, pLink, ProteinProspector, and Kojak use the top 500, 1000, and 250 peptides, respectively, to generate peptide-peptide pairs. Such a strategy, however, only searches a fraction of all possible peptide-peptide pairs. Let’s take the Homo sapiens (human) database as an example. There are around 10^7^ peptide-peptide pairs for each experimental spectrum. With a rough estimation, pLink, ProteinProspector, and Kojak only search about 1.2% 5.0%, and 0.3% of all peptide-peptide pairs, respectively. Clearly, the non-exhaustive search strategy cannot guarantee to find a spectrum’s highest scored peptide-peptide pair. The result is highly variable with respect to the database size. Our experiments have shown that there is a significant proportion of pairs missing with the non-exhaustive search strategy^37^. Also, the sensitivity decreases greatly as the size of the database increases. Petrotchenko and Borchers ^38^ proposed a fast algorithm to search peptide-peptide pairs. Unfortunately, that algorithm’s time complexity is quadratic in theory and it has already been widely used in tools including xQuest, Kojak, and ECL. We will show that the time complexity is quadratic using a large-scale data set.

In XL-MS, using a noncleavable amine-reactive cross-linker (such as DSS and BS3) to link two proteins is a common protocol^23,35,36,39,40^. In order to process XL-MS data using a noncleavable cross-linker, we developed a tool named ECL^37^ that can exhaustively search a database in dozens of hours. However, the running time of ECL increases quadratically with respect to the database size. Comparison with benchmark methods in the second group shows that ECL is faster than xQuest, pLink, and StavroX, but is slower than ProteinProspector ^25^ and Kojak^27^ when the database is large. Furthermore there is a high possibility that a sample contains a large number of proteins. Such phenomenon is very common in *in vivo* experiments. Take the data set published by Zhu et al.^6^ for example, there are in total 15976 proteins that possibly exist in the sample. This mean that the database used for cross-linked peptides identification must be large. Unfortunately, for those that can handle a large database, they rely on a non-exhaustive search strategy which results in a significant proportion of missing findings^37^. Thus, it is desirable to further speed up the computation of ECL.

In this paper, we propose a tool named ECL 2.0 that can search all peptide-peptide pairs with a linear time and space complexity. ECL 2.0 has several advantages over ECL: achieving a linear time complexity; using XCorr^41,42^as the score function, which is more robust to noise peaks than the correlation coefficient used in ECL; using a linear-tail-fit method^43^ to estimate *e*-value for each PSM, which statistically calibrates the original score function. ECL 2.0 can analyze a typical LC-MS/MS data set using a large-scale database containing thousands of proteins in a few hours. To our knowledge, no exhaustive search tool can achieve such a speed. ECL 2.0 also identifies the largest number of PSMs compared to pLiuk. StavroX. ProteiuProspector. kojak. and ECL when using the same false discovery rate (FDR)/*q*-value threshold.

The rest of the paper is organized as follows: Section 2 describe the algorithm of ECL 2.0. Section 3 demonstrates the performance of ECL 2.0 with real data sets. Section 4 concludes the paper with some discussions

## 2 Method

By convention^10,23^, two linked peptides are called α chain and *β* chain, respectively. After dissociation (e.g. CID), there are fragmented ions from two peptide chains. Figure 1 illustrates the ions with marks: green marks indicate linear ions that only contain one chain’s ions; red marks indicate cross-linking ions that contain one chain’s ions plus a modification containing the cross-linker and the other whole peptide.

**Figure 1:**
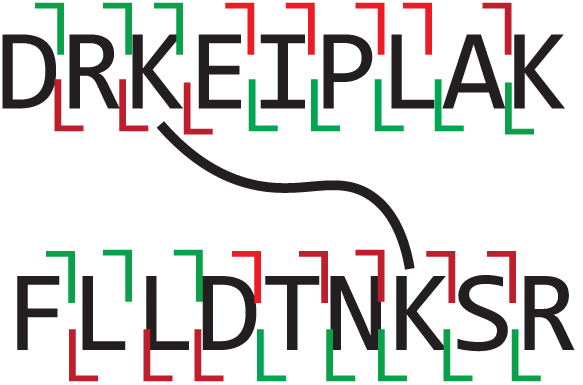
An illustration of cross-linked peptides. Green marks indicate linear ions and red marks indicate cross-linking ions.

Given an experimental spectrum, the objective of cross-linked peptides identification can be expressed as

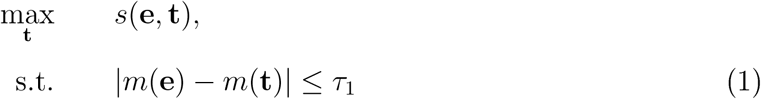

where *s*(**e, t**) is a score function, **e** is an experimental spectrum, **t** is a theoretical spectrum of cross-linked peptides, *m*(**•**) is the precursor mass of a spectrum, and *τ*_1_ is the precursor mass tolerance. We need to pair two peptide chains to generate the corresponding theoretical spectra, which results in a quadratic time complexity.

With an additive score function^2,44^, the score corresponding to a peptide-peptide pair equals the sum of two scores corresponding to two peptide chains. Other researchers have also made such observation^17,21,27^, but they did not take advantage of it to reduce the time complexity. Based on this observation, we propose a new algorithm that achieves a linear time complexity. Our algorithm can be applied to any score function with an additive property.

### 2.1 Additive Score Function

Given a spectrum, we can digitize the whole *m/z* range into bins based on the MS2 *m/z* tolerance:

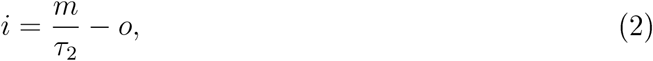

where *i* is the index of the digitized bin, *m* is an *m/z* value, *τ_2_* is the MS2 *m/z* tolerance, and *o* is an offset. For the *i*-th bin, the corresponding intensity can be obtained as

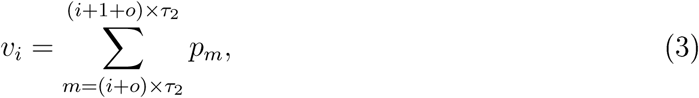

where *υ_i_* is the *i*-th value in the digitized vector and *p_m_* is the peak intensity whose *m/z* value is *m*. If there is no peak at the location *m, p_m_* = 0. Then, we have the following definition:

#### Definition 1

*Given a digitized experimental spectrum e and a digitized theoretical spectrum* t*, an additive score function reads*

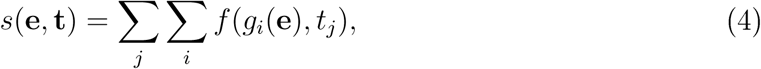

*where g_i_*(*e*) *is a measure of the i-th bin in the experimental spectrum, t_j_ is the j-th value in* **t**, *and f* (*g*_i_(*e*), *t_j_*) *is a score term.*

Roughly speaking, there are two types of score functions in linear/cross-linked peptides identification tools^21,23,25,27,35,36,41,45–49^:

1. Dot-product-based score functions, such as the dot product^25,47,48^, intensity summation^35,36^, XCorr^21,27,35,36,11,15^. and the kernel spectral dot product (KSDP) ^23,46^.
2. Probability based score functions, such as the match-odds^35,36^ and log-odds function^49^.

The dot product, intensity summation and XCorr are additive score functions. For example, XCorr can be expressed as:

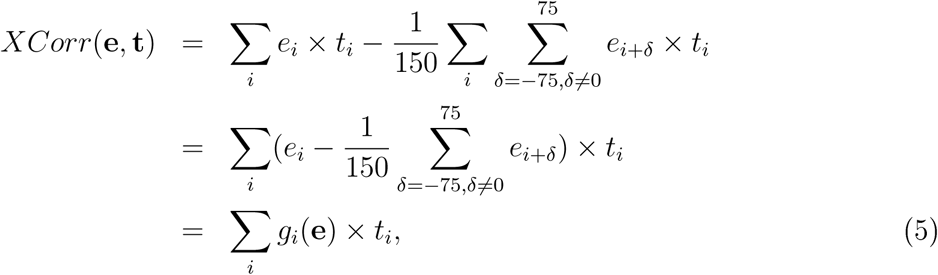

where *δ* is an *m/z* offset and 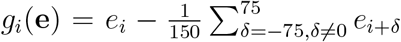 Here, we assume that there are no overlapping peaks in the theoretical spectrum. This assumption may not be true in some cases. However, such cases are quite rare due to the high resolution property of mass spectrometer (e.g. Thermo Scientific Q-Exactive and LTQ Orbitrap Elite).

According to Figure 1, theoretical peaks are from four sources: linear ions and crosslinking ions from α chain; linear ions and cross-linking ions from *β* chain. Correspondingly, a score of two linked peptide chains can be expressed as

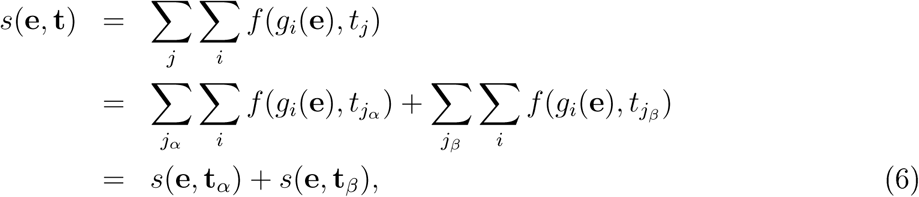

where *j*_*α*_ is a bin index corresponding to the peak from α chain, *j_β_* is a bin index corresponding to the peak from *β* chain, t_**α**_ is a digitized theoretical spectrum containing peaks from α chain only, and t_***β***_ is a digitized theoretical spectrum containing peaks from *β* chain only. Let’s call s(e, t_**α**_) and *s*(**e**, *t_β_*) chain scores for convenience. With Equation (6), Equation (1) can be expressed as

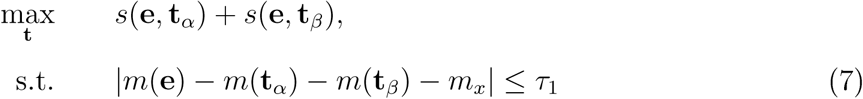

where *m*_***x***_ is the mass of the cross-linker.

### 2.2 Searching Cross-Linked Peptides

Equation (7) implies that, given an experimental spectrum, we can first calculate all chain scores separately (Algorithm 1). Then, we can add pairs of chain scores. The time complexity of adding all possible pairs of scores is quadratic with respect to the number of chain scores. Here, we propose a digitization-based algorithm to achieve a linear time complexity. We first describe the procedure of searching cross-linked peptides given an experimental spectrum. Then, we analyze this procedure’s time and space complexity.

#### Algorithm 1

Calculating chain scores.

Without loss of generality, we use ions fragmented from CID and don’t consider neutral loss ions. It can be easily applied to other dissociation methods by changing b/y-ion to a/x-ion or c/z-ion. When we consider neutral loss ions, the computational complexity does not change, **b** is a vector of **b**-ion masses from the peptide chain, and **y** is a vector of y-ion masses from the peptide chain. We assume that the mass difference of ions is larger than the MS2 *m/z* tolerance, *x* is the mass of the cross-linker, *m_c_* is the mass of the peptide chain, e is the digitized experimental spectrum, *m_e_* is the mass of the experimental spectrum, *τ_2_* is the MS2 *m/z* tolerance, and *o* is the offset in digitization.

~~~
1: **procedure** CHAINSCORE(**b**, **y**, *x, m_c_*, **e**, *m_e_*, τ_2_, *o*)
2:   c*← υector*[*len*(e)]
3:   *s ←* 0
4:
5:   **for** *i ←* 1*; len*(**b**) do            ⊳ fill bins corresponding to b-ions
6:      **if** *i < x* **then**
7:      **c**[⌊*b_i_/T*_2_ + *o*⌋] *←*1 ⊳ ⌊*b_i_/T_2_ + o*⌋ is a function taking the largest integer smaller than *b_i_/T_2_* + *o*
8:   **else**
9:      **c**[⌊(*b_i_* + *m_e_ – m_c_*)*/T*_2_+ *o*⌋] *←* 1
10:    **end if**
11:  **end for**
12:
13:   for *i ←* 1*; len*(y) **do** ⊳ fill bins corresponding to y-ions
14:      **if** *i > x* then
15:     **c** [⌊*y_i_/T*_2_ + *o*⌋] *←* 1
16:      **else**
17:     **c**[⌊(*y_i_* + *m_e_ – m_c_*)/τ_2_+*o*⌋] *←* 1
18:      **end if**
19:    **end for**
20:
21:    **for** *j ←* 1*; len*(c) **do** ⊳ calculate the chain score
22:       **for** *i ←* 1*; len*(e) **do**
23:           *s ← s* + *f*(*g_i_*(e); c[*j*]) ⊳ *f*(*g_i_*(e); c[*j*]) is the score function
24:      **end for**
25:     **end for**
26:
27:    **return** *s*
28: **end procedure**
~~~

Given a database, we first *in silicon* digest all proteins into *n* peptide chains. All the peptide chains are sorted based on their masses. Then, we split the whole mass range into multiple intervals. The width of the intervals *w* is much smaller than the precursor mass tolerance *τ*_1_ Here, we set the width *w* to 0.001 Da. The number of intervals is equal to

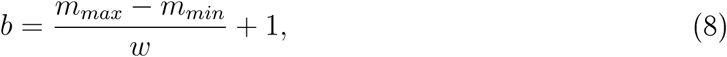

where *b* is the number of intervals, *m*_***max***_ is the maximal mass of peptide chain, and *m*_***min***_ is the minimal mass of peptide chain. All of these values are pre-fixed before the database search. Finally, we assign peptide chains into different intervals based on their masses, and all peptide chains in the same interval are treated as having the same mass.

Given an experimental spectrum, we calculate the chain scores with respect to all possible peptide chains using Algorithm 1 and assign them to the corresponding intervals. According to Section 2.1, the highest score must come from one of the following situations:

- Two peptide chains are from different intervals: the highest score is equal to the sum of the two top chain scores in two different intervals.
- Two peptide chains are from the same interval: the highest score is equal to two times the top chain score in the interval.

Thus, we only need to keep the top-scored peptide chain and the chain score in each interval during the calculation of *s*(e, t). Given a peptide chain in the *i*-th interval, another peptide chain must be in the interval satisfying

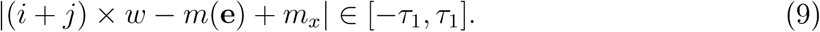

Here, we ignore the rounding error because *w* (i.e. 0.001 Da) is much smaller than *τ*_1_ (e.g. 0.05 Da) in practice. Algorithm 2 shows the pseudo code of identifying cross-linked peptides given a set of peptide chains and an experimental spectrum.

#### Algorithm 2

Identifying cross-linked peptides with a linear time and space complexity. Without loss of generality, we use ions fragmented from CID and don’t consider neutral loss ions. {*b_i_*} is a set of b-ion mass vectors from all peptide chains, {y_*i*_} is a set of y-ion mass vectors from all peptide chains, {*x_i_*} is a set of link-site indexed corresponding to all peptide chains, {*m_ci_*} is a set of peptide chain masses, e is a digitized experimental spectrum, *m_e_* is the mass of the experimental spectrum, *m_x_* is the mass of the cross-linker, τ_1_ is the precursor mass tolerance, τ _2_ is the MS2 *m/z* tolerance, and *o* is the offset in digitization.

~~~
1: **procedure** SEARCH({b_*i*_},{y_*i*_},{*x_i_*},{*m_ci_*}, e, *m_e_*, *m_x_*, τ_1_, τ_2_,o) ⊳ *i* ∈[1,*n*]
2:   *s_c_ ← υector*[⌈max({*m_Ci_*})*/* τ_1_⌉]
3:   *s ←* 0 ⊳ *s* is the final score
4:   *c*_1_ *← –*1 ⊳ *c*_1_ is the index of the first peptide chain in the final result
5:   *c*_2_ *← –*1 ⊳ *c*_2_ is the index of the second peptide chain in the final result
6:
7:   **for** *i ←* 1*; |*{*m_ci_}|*do ⊳ calculate chain scores and assign them to ranges
8:      *s ←* ChainScore(b*_i_*, y*_i_, x_i_;m_ci_*, e, *m_e_*, τ_2_,*o*)
9:      **if** s_*c*_[⌊*m_ci_/T*_1_⌋] *< s* **then**
10:        **s**_*c*_ [⌊*mc_i_ / T*_1_⌋] *← s*
11:    **end if**
12:   **end for**
13:
14:   **for** *i ←*1, ⌈max({*m_ci_*})*/*0:001⌉*/*2 **do** ⊳ pair peptide peptide pairs
15:      **for** *j ←* (*m_e_ – m_x_ – m_ci_ –* τ_1_)/*=*0:001 *– i*, (*m_e_ – m_x_ – m_ci_* + τ_1_)*/* 0:001 *– i* **do**
16:        **if** s_*c*_[*i*] + _s*c*_[*j*] *> s* **then**
17:            *s ←* **s***c*[*i*] + **s***c*[*j*]
18:            *c*_1_ *← i*
19:            *c*_2_ *←j*
20:         **end if**
21:       **end for**
22:    **end for**
23:
24:   **return** *s; c*_1_*,c*_2_
25: **end procedure**
~~~

In the following, we analyze the time and space complexity of Algorithm 2. The time complexity of mass range splitting and peptide chains assignment is

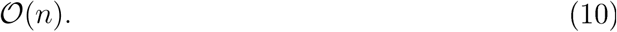

Without loss of generality, we suppose that the time and space complexity of calculating a chain score is independent of the number of peptides. The time complexity of calculating all chain scores and assigning them to intervals is

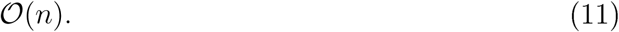

The time complexity of finding pairs of peptide chains, summing chain scores, and keeping the highest-scored pair is

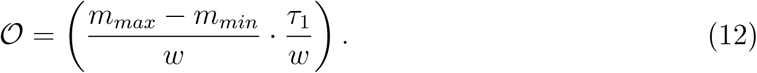

Combining Equation (10), (11), and (12), we obtain the total time complexity:

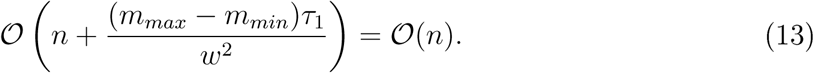

*Because m_max_*, *m_min_*, *T_1_ and w are pre-fixed and independent of the database size, the total time complexity is linear with respect to the number of peptide chains. It is easy to see that the space complexity is also linear:* 𝒪 (*n* + (*m_max_* – *m_min_*)/*w*).

In summary, the linear time complexity is achieved with the following factors:

1) Taking advantage of an additive score function: A final score can be split into two additive chain scores.
2) Digitizing the whole mass range and assigning peptide chains to digitized intervals: With such a digitization, only one score is kept for each interval. This reduces the time complexity greatly. Given a peptide chain, the time complexity of finding another peptide chain having the highest chain score is 𝒪 (n log(*n*)) with the help of quicksort^50^. Thus, the bottleneck lies in sorting.
3) Achieving a constant time complexity in summing up chain scores by fixing the number of digitized intervals. By fixing the number of ranges, we eliminate the bottleneck and the total time complexity reduces to 𝒪 (*n*). Actually, using counting sort can achieve 𝒪 (*n*) time complexity^50^ without fixing the number of ranges. However, its space complexity is quite high, which will be an issue in cross-linked peptides identification.

### 2.3 The Work-Flow of ECL 2.0

The major contribution of this paper is proposing a linear computational complexity algorithm to exhaustively searching all peptide-peptide pairs in cross-linked peptides identification task. Since XCorr has been widely used ^27,11,15,51^ and has a relatively high sensitivity among other score functions^46,52,53^, we choose it as the score function of ECL 2.0. Figure 2 shows the work-flow of ECL 2.0. It takes a data file and a protein database file as inputs. After digitizing spectra, digesting protein sequences, and *in silicon* fragmenting peptide sequences, it calculates chain scores using Algorithm 1. Once is has obtained each spectrum’s chain scores, the method pairs peptide chains and finds the highest-scored pair using Algorithm 2. It then calculates an e-value using the linear tail-fit method^43^.

**Figure 2:**
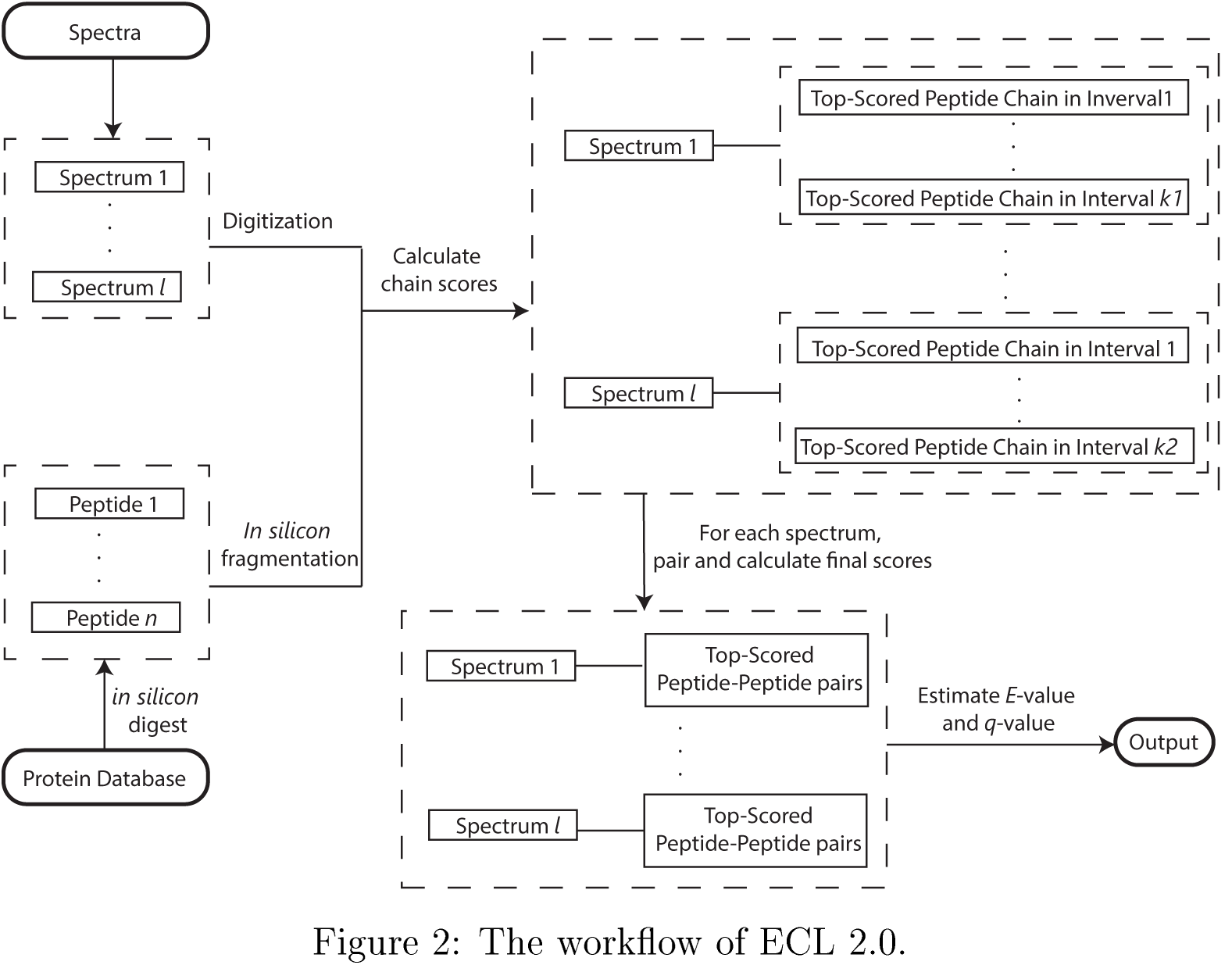
The workflow of ECL 2.0.

There are three kinds of PSM: the first contains two peptide chains from the target database, the second contains two peptide chains from the decoy database, and the third contains one peptide chain from the target database and another peptide chain from the decoy database. Thus, we can estimate the FDR using^23,36^

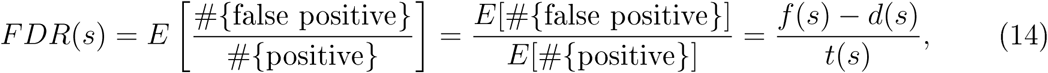

where s is an *e*-value thresh old, #{false positive} is the number of false positives with *e-*values equal to or smaller than *s*, #{positive} is the number of positives with *e*-values equal to or smaller than *s, E*[•] is the expectation, *t*(*s*) is the number of the first kind of PSMs with *e*-values equal to or smaller than *s, d*(*s*) is the number of the second kind of PSMs with *e*-values equal to or smaller than s, and *f* (*s*) is the number of the third kind of PSMs with *e*-values equal to or smaller than *s.* In order to make the sensitivity as high as possible, FDR is converted into *q*-values in proteomics tools, such as crux^51^, Comet^45^, Percolator^54^, pLink^23^, xProphet^36^, and ECL^37^. In ECL 2.0, we also do such a conversion using^55^

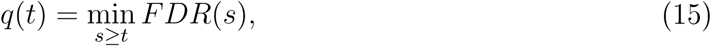

where *t* is a threshold.

## 3 Experiments

We perform two sets of experiments to show the power of ECL 2.0. The first set contains 10 data files in Makowski et al. ^56^. The purpose is to demonstrate the effect of database size and the performance of ECL 2.0 under different settings. The authors published 19 confident cross-linked proteins. We use them to generate databases with six different sizes. We use pLink, StarvoX, ProteinProspector, Kojak, and ECL as benchmarks. The second set contains 30 data files in Zhu et al.^6^. The data files are from *in vivo* cross-linked proteins of *Arabidopsis thaliana.* The whole *Arabidopsis thaliana* database contains more than 3.5 × 10^4^ proteins^57^. All the detailed parameter settings, result hies, and logs are available at Supplementary File.

### 3.1 Identifying Peptides from *In Vivo* Cross-Linked *Homo Sapiens* Proteins

In Makowski et al.^56^, there are 10 data files from the cross-linking of human proteins, containing about 3 × 10^5^ MS2 spectra in total. Please refer to Makowski et al.^56^ for details of the sample preparation and data acquisition. The authors reported 19 cross-linked proteins with a high conhdence. In real applications, the database usually contains more proteins that do not exist in the sample. In the following, we combine different numbers of additional proteins with the 19 proteins to generate six databases:

- The first database contains the 19 proteins plus 50 randomly selected proteins.
- The second database contains all proteins in the first database plus another 150 randomly selected proteins.
- The third database contains all proteins in the second database plus another 800 randomly selected proteins.
- The fourth database contains all proteins in the third database plus another 4000 randomly selected proteins.
- The fifth database contains all proteins in the fourth database plus another 5000 randomly selected proteins.
- The last database contains all proteins in the fifth database plus another 5000 randomly selected proteins.

The randomly selected proteins are from *Arabidopsis thaliana*, while the samples are from *Homo sapiens.* Without considering decoy sequences, we have six databases whose protein numbers are 69, 219, 1019, 5019, 10019, and 15019, respectively.

We use StavroX (Version 3.6.0), pLink (Version 1.23), ProteinProspector (Version 5.17.1), Kojak (Version 1.5.3), ECL (Version 1.1.1), and ECL 2.0 (Version 2.1.2) to search these data hies against six databases, respectively. The precursor mass tolerance is 10 ppm, and the MS2 *m/z* tolerance is 0.01 Da. The allowed maximum missed cleavage is two. The allowed precursor masses are from 1000 Da to 12000 Da and the allowed peptide chain lengths are from 5 amino acids to 50 amino acids. We set carbamidomethylation on “C” as the fixed modification. We don’t set any variable modification. All six tools use the target-decoy strategy^23,36,58^ to estimate the FDR and *q-*value. The decoy sequences are generated by reversing target sequences with C-terminal unchanged. We use *q*-value ≤ 0.05 as the threshold for these tools. StavroX and pLink provide q-values for their own results. We use Percolator^54^ to estimate *q*-values for Kojak’s results, as advised by the authors^59^. ProteinProspector doesn’t provide *q*-value in the result. Thus, we estimate it using Equation (14) and Equation (15). ECL and ECL 2.0 report *q*-values by themselves. With the cut-offed results, the PSMs from the cross-linking of the 19 proteins are treated as true positive PSMs and the PSMs containing at least one peptide chain from the randomly selected proteins are treated as false positive PSMs.

#### 3.1.1 Identification Results

We summarize the true positive PSMs and false positive PSMs identified by six tools. We find that StavroX and Kojak may report multiple peptide-peptide pairs for one spectrum. For fair comparison, we only count each of these spectra once. Figure 3 shows the bar plots of true positive and false positive PSMs. StavroX cannot handle the second to sixth databases, pLink cannot handle the fourth to sixth databases, ProteinProspector cannot handle the fifth and sixth databases, and ECL needs many days to search the fifth and sixth databases.

**Figure 3:**
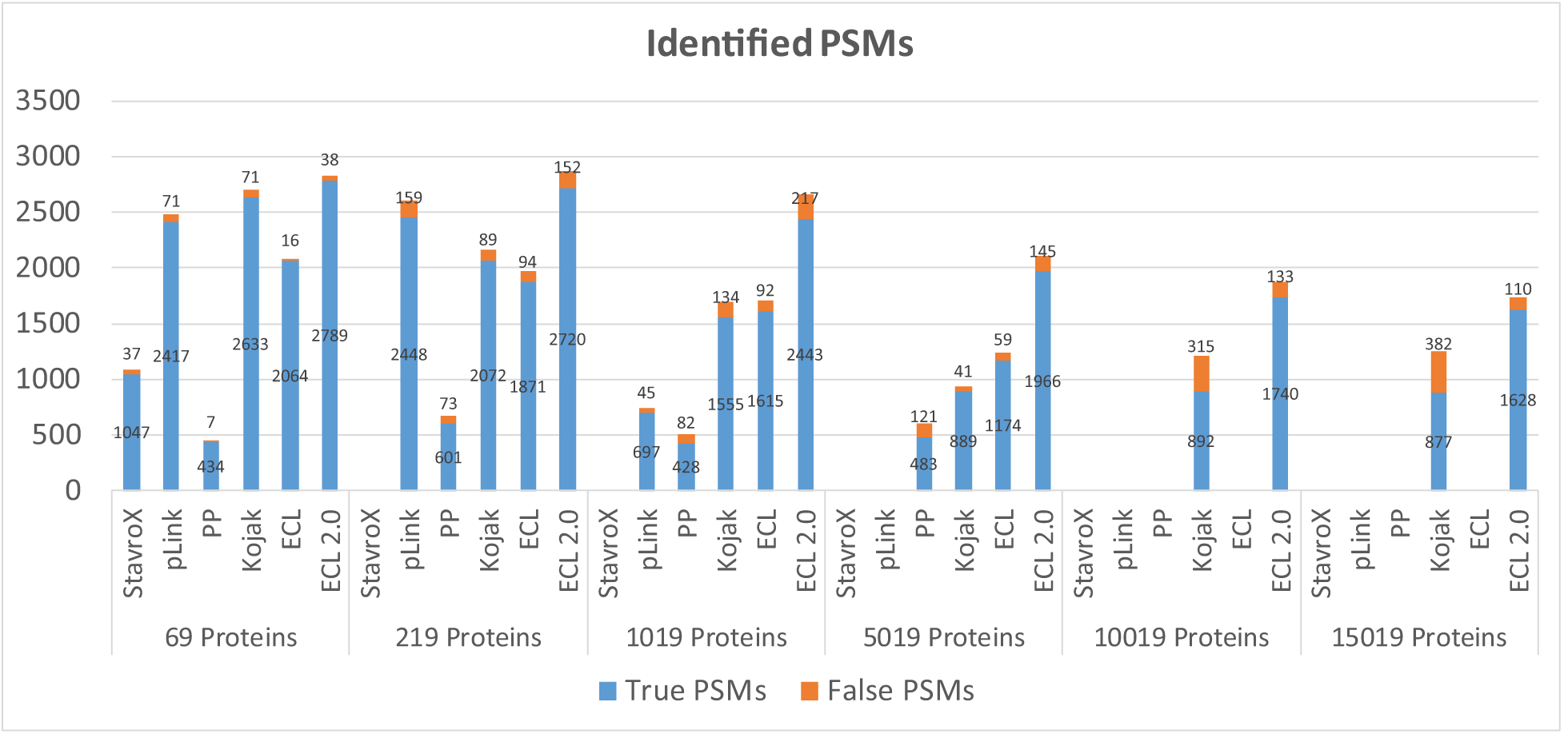
Bar plots showing identified true and false positive PSMs. Six bar plots correspond to the results of searching six databases. Without considering decoy proteins, there are 69, 219, 1019, 5019, 10019, and 15019 proteins in six databases, respectively. The blue bars denote true positive PSMs and the orange bars denote false positive PSMs. The value in the middle of each blue bar is the number of corresponding true positive PSMs and the value at the top of each orange bar is the number of corresponding false positive PSMs. “PP” stands for ProteinProspector. StavroX cannot handle the second to sixth databases, pLink cannot handle the fourth to sixth databases, ProteinProspector cannot handle the fifth and sixth databases, and ECL needs many days to search the fifth and sixth databases. Thus, the corresponding bars are left blank.

Thus, the corresponding bars are left blank. The blue bars denote the true positive PSMs and the orange bars denote the false positive PSMs. The value in the middle of each blue bar is the number of corresponding true positive PSMs and the value at the top of each orange bar is the number of corresponding false positive PSMs.

Figure 3 shows that ECL 2.0 always identifies the largest number of PSMs among all six tools. Although both Kojak and ECL 2.0 use XCorr as the score function, Kojak reduces the computational burden with four simplifications ^59^:

- Using at most 250 peptide chains to generate peptide-peptide pairs for each experimental spectrum.
- Using a so called “turbo_button” function to reduce the number of searched peptides furthermore.
- Removing small value peaks (from −0.5 to 0.5) from preprocessed experimental spectra.
- Using the char data type (one byte with integer ranges from −128 to 127) rather than the float data type (four bytes with floating point values) to represent the preprocessed experimental spectra.

ECL 2.0 doesn’t simplify anything. Furthermore, ECL 2.0 also estimates *e*-value for each PSM to calibrate the original score. ECL uses a score function different from that of ECL 2.0 and it doesn’t calibrate the score with *e*-value. Thus, it identifies a smaller number of PSMs.

It is also interesting that as the database size increases, the number of identified PSMs decreases for all tools. One reason is that the chance of random matching increases as database size increases. In order to maintain the same *q*-value, a higher score threshold is required. Thus, the number of PSMs passing the corresponding score threshold decreases. All six tools suffer from this reason. The other reason lays in the non-exhaustive search strategy. As the increase of database size, the chance of missing the underlying true PSMs increases (Figure S3). Tools except for ECL and ECL 2.0 suffer from this reason. Figure 3 shows that ECL and ECL 2.0 are less affected.

#### 3.1.2 Running Time

In this section, we calculate each tool’s average running time with respect to different database sizes. StavroX, pLink, Kojak, ECL, and ECL 2.0 are run on a PC with an Intel Core i7-2600 CPU (3.40 GHz, 8 cores) and 32 GB memory. StavroX, pLink, and ECL don’t support multi-thread computing. Kojak and ECL 2.0 support multi-thread computing. Thus, Kojak and ECL 2.0 are run with 8 cores. ProteinProspector is run on the authors’ web server. Table 1 shows the average running time in hours. ECL 2.0 is much faster than StavroX, pLink, ProteinProspector, and ECL. But it is slower than Kojak. We have explain earlier that Kojak reduces the computational burden with four simplifications. We argue that the simplifications are the major reasons making Kojak faster than ECL 2.0.

**Table 1:** The average running time of six tools with respect to different database sizes. The unit is hour. StavroX cannot handle the second to sixth databases, pLink cannot handle the fourth to sixth databases, ProteinProspector cannot handle the fifth and sixth databases, and ECL needs many days to search the fifth and sixth databases. Thus, the corresponding cells are marked with “ΝΑ”. “PP” stands for ProteinProspector.

|  | Target Protein Number in the Database |  |  |  |  |  |
| --- | --- | --- | --- | --- | --- | --- |
|  | 69 | 219 | 1019 | 5019 | 10019 | 15019 |
| StavroX | 5.11 | NA | NA | NA | NA | NA |
| pLink | 30.25 | 10.40 | 33.62 | NA | NA | NA |
| PP | 0.74 | 1.52 | 1.32 | 7.77 | NA | NA |
| Kojak | 0.04 | 0.06 | 0.11 | 0.43 | 0.75 | 1.13 |
| ECL | 0.10 | 0.61 | 5.35 | 85.44 | NA | NA |
| ECL 2.0 | 0.21 | 0.33 | 0.88 | 3.61 | 8.33 | 12.14 |

In order to show that ECL 2.0 does have a linear time complexity, we plot the average running time with respect to the numbers of peptide chains (including decoy sequences) in Figure 4. For comparison, we also plot the average running time of ECL (version 1.1.1) that has a quadratic time complexity. The figure shows the advantage of linear complexity clearly.

**Figure 4:**
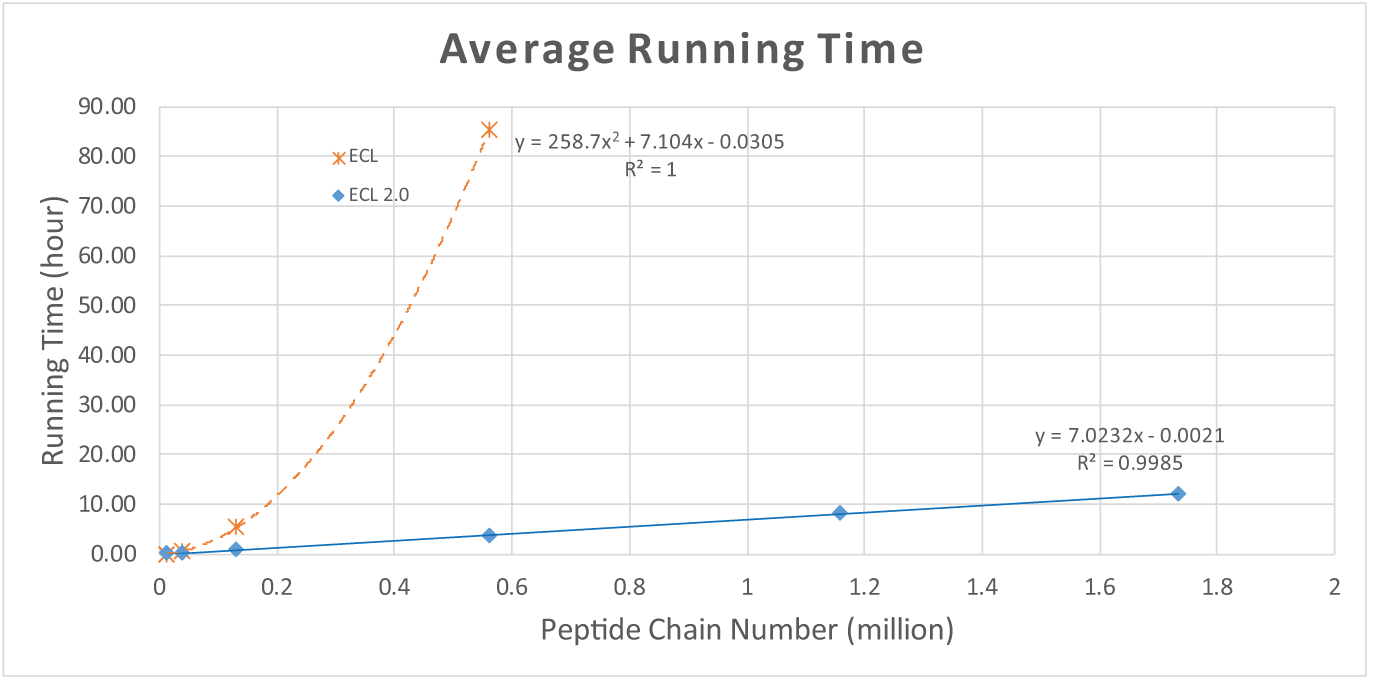
A plot showing average running time with respect to different numbers of peptide chains. Decoy sequences are included. The *x*-axis is the number of peptide chains (the unit is million) and the *y*-axis is the average running time (the unit is hour). The orange crosses are the observed running time of ECL and the orange dashed line is the fitted quadratic line. The blue dots are the observed running time of ECL 2.0 and the blue solid line is the fitted linear line. It also shows two lines’ equations and R^2^ values.

**Figure 5:**
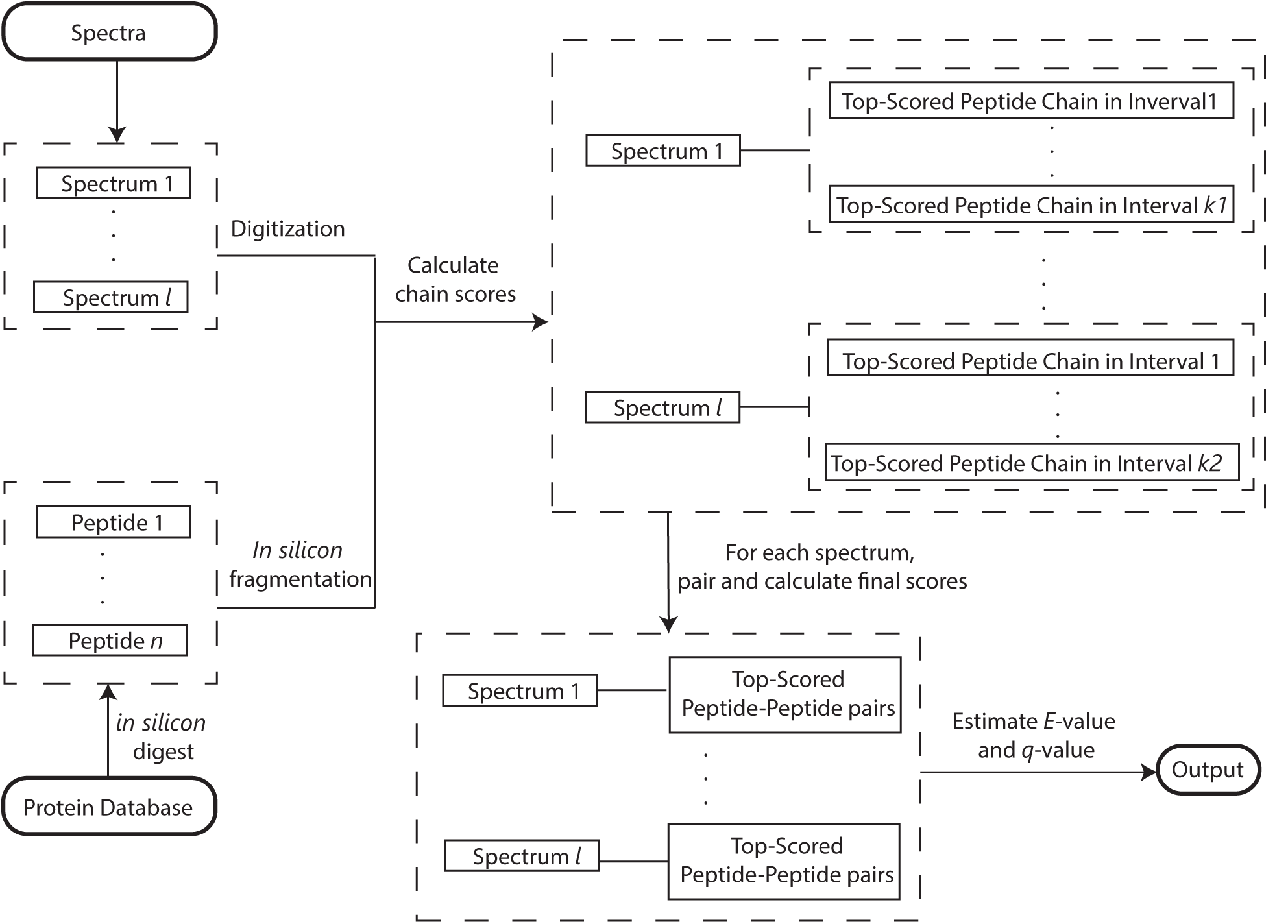
For TOC only.

### 3.2 Identifying Peptides from *In Vivo* Cross-Linked *Arabidopsis Thaliana* Proteins

In this section, we use a large-scale cross-linking data set to demonstrate the power and necessity of ECL 2.0. 30 data files from *in vivo* cross-linked *Arabidopsis thaliana* proteins are collected in Zhu et al. ^6^. There are around 6 × 10^5^ MS2 spectra in total. Please refer to Zhu et al.^6^ for the details of sample preparation and data acquisition. Since the data is from *in vivo* cross-linking and we don’t have the ground truth, we use the whole proteome of *Arabidopsis thaliana* as the database (download from the Arabidopsis information resource^57^). There are more than 3.5 × 10^4^ proteins. Considering oxidation as the variable modification, there are more than 2 × 10^6^ peptides which form 2 × 10^12^ peptide-peptide pairs.

We try StavroX (Version 3.6.0). pLink (Version 1.23), ProteinProspector (Version 5.17.1), kojak (Version 1.5.3), ECL (Version 1.1.1), and ECL 2.0 (Version 2.1.2) to identify this data set. Because the data is generated by Thermo LTQ Orbitrap XL mass spectrometer, the precursor mass tolerance is 10 ppm and the MS2 *m/z* tolerance is 0.5 Da. We set the allowed maximum missed cleavage to one. We set carbamidomethylation on “C” as the fixed modification and oxidation on “M” as the variable modification. Rest of the parameters are the same as the earlier experiment.

Because the database is too large for a PC, we run the tasks on a server with two Intel Xeon E5-2670 v3 CUPs (24 cores in total) and 128 GB memory. Unfortunately, only kojak and ECL 2.0 can finish the task within a reasonable period of time. For each data file. Kojak needs about one hour and ECL 2.0 needs about five hours, respectively. With *q*-value ≤ 0.05 as the threshold, Kojak doesn’t identify any PSM but ECL 2.0 identifies 96 PSMs including those two reported by Zhu et al. ^6^. To take a closer look, we summarize the ranks of peptide chains corresponding to each spectrum using ECL 2.0. It turns out that 77 out of 95 PSMs have at least one peptide chain whose rank is lower than 250. Since Kojak uses at most 250 peptide chains to generate peptide-peptide pairs for each spectrum, those PSMs will be missed. We also notice that 54 out of 95 PSMs have at least one peptide chain whose rank is lower than 1000. This means that even setting peptide chain number to that of ProteinProspector, there are still significant missing findings. The detailed results can be found in Table S1.

## 4 Discussions

In this paper, we demonstrated that it is feasible to exhaustively search all possible peptide-peptide pairs with a linear time and space complexity. Given a data file with tens of thousands of MS2 spectra, ECL 2.0 can finish the analysis using a big database in a few hours. Our experiments showed that the missing findings caused by non-exhaustive search become more and more critical as the increase of database size. When the database becomes as huge as containing more than 3.5 × 10^4^ proteins, only ECL 2.0 can find PSMs passing the FDR threshold. This demonstrates the power of exhaustive search. Since, most biological experiments involving many proteins, exhaustive search is necessary and will be useful in analyzing most cross-linking data.

For each MS/MS spectrum, ECL 2.0 has to calculate chain scores corresponding to peptides in a wide precursor mass range, which takes most of the computational time. From our point of view, this is the only issue left in exhaustively identifying cross-linked peptides. We will try to address this issue in the future.

## Abbreviations

ECL: exhaustive cross-linked peptides identification.
XL-MS: chemical cross-linking coupled with mass spectrometry.
ppm: parts per million.
CID: collision-induced dissociation.
MS1: first level mass spectrometry.
MS2: tandem mass spectrometry.
MS3: third level mass spectrometry following MS2.
DSS: disuccinimidyl suberate.
BS3: bis(sulfosuccinimidyl) suberate.
KSDP: kernel spectral dot product.
FDR: false discovery rate.
PSM: peptide-spectrum match.

## Acknowledgement

This work is partially supported by a theme-based project T12-402/13N from the Research Grants Council (RGC) of the Hong Kong S.A.R. government, an internal grant VPRGO15EG01 from HKUST, grants 16101114 and 661613 from the General Research Fund (GRF) of the Hong Kong S.A.R. government, and grant 31370315 from the National Natural Science Foundation of China (NSFC). We thank Meng Wang and Jiaan Dai for valuable discussions.

## Supporting Information

This information is available free of charge at http://pubs.acs.org/

- Supplementary Document. (PDF)
- Supplementary Table S1. (XLSX)
- Supplementary File: protein databases, all tools’ parameter files, result files, and logs. (ZIP)

The source code and executable file can be found at http://bioinformatics.ust.hk/ecl2.0.html.

